# Contrasting patterns of *Asaia* sp association with *Plasmodium falciparum* between field collected *Anopheles gambiae* and *Anopheles coluzzii* from Cameroon

**DOI:** 10.1101/2024.03.04.583377

**Authors:** Claudine Grâce Tatsinkou Maffo, Maurice Marcel Sandeu, Micareme Tchoupo, Fleuriane Metissa Dondji Kamga, Leon M.J. Mugenzi, Flobert Njiokou, Grant L. Hughes, Charles S. Wondji

**Affiliations:** Department of Medical Entomology, Centre for Research in Infectious Diseases (CRID), LSTM Research Unit, P.O. BOX 13591, Yaoundé, Cameroon; Department of Animal Biology and Physiology, Faculty of Science, University of Yaoundé 1, P.O. Box 812, Yaoundé, Cameroon; Department of Microbiology and Infectious Diseases, School of Veterinary Medicine and Sciences, University of Ngaoundéré, Po Box 454, Ngaoundéré, Cameroon; Departments of Vector Biology and Tropical Disease Biology, Centre for Neglected Tropical Diseases, Liverpool School of Tropical Medicine, Liverpool, United Kingdom; Department of Vector Biology, Liverpool School of Tropical Medicine, Pembroke Place, Liverpool, United Kingdom

**Author notes:** **Corresponding author:** Maurice Marcel SANDEU, Centre for Research in Infectious Diseases (CRID), 39 Rue Marcus Etoundi P.O. Box 13591, Yaoundé Cameroon. **Authors’ information** Claudine Grâce Tatsinkou Maffo and Maurice Marcel Sandeu contributed equally to this work and share first authorship.

**Keywords:** Asaia, *Anopheles gambiae*, *Anopheles coluzzii*, *Plasmodium falciparum*, Seasons

## Abstract

The presence of bacteria of the genus *Asaia* in mosquitoes makes them suitable candidates for malaria paratrangenic control. To better understand whether this bacterium could be used for malaria control, we investigated and quantified *Asaia* influence in *An.*L*gambiae sl* populations naturally infected by the malaria parasite *Plasmodium falciparum.* Adult *Anopheles* mosquitoes were collected across two different eco-geographical localities in Cameroon, during the dry and wet seasons. DNA was extracted from the whole individual mosquitoes and *Asaia* was quantified using real-time Q-PCR by amplification of 16S ribosomal RNA gene. We also detected and quantified *Plasmodium falciparum* infection in same mosquitoes. The density of *Asaia* was quantified in 864 mosquitoes. This included 439 *An. gambiae* collected from Bankeng and 425 *An. coluzzii* collected from Gounougou. The prevalence of *Asaia* in *An. gambiae* and *An. coluzzii* was 88.3% and 80.9% respectively, with *An. gambiae* were more heavily infected by *Asaia* than *An. coluzzii.* There was also a significant difference between the infection densities of *Asaia* per season in the two localities. We also had a significant difference between the infection densities of *Asaia* according to *Plasmodium* infection status in the two localities. However, no correlation was observed between the amount of *Asaia* and *P. falciparum* infection. This study provides evidence that naturally occurring *Asaia* infection is not correlated to *P. falciparum* development within *An. gambiae* and *An. coluzzii.* Further studies as experimental infection and innate immune response are required to better investigate the correlation between *Anopheles* mosquitoes, *Asaia* and *Plasmodium*.

**Importance:** Malaria control relies mainly on insecticide-based tools. However, the effectiveness of these tools is threatened by the widespread insecticide resistance in malaria vectors highlighting the need for alternative control approaches. The endosymbiont *Asaia* has emerged as a promising candidate for paratransgenic control of malaria, but its biology and genetics still need to be further analyzed across Africa. Here, we investigated and quantified *Asaia* influence in *An.*L*gambiae sl* populations naturally infected by the malaria parasite *Plasmodium falciparum.* DNA was extracted from the whole individual mosquitoes collected from two localities, and then *Asaia* was quantified using real-time Q-PCR by amplification of 16S ribosomal RNA gene. We also detected and quantified *Plasmodium falciparum* infection in same mosquitoes and correlated *Plasmodium* infection them with the presence of *Asaia* infection. This study provides evidence that naturally occurring *Asaia* infection is not correlated to *P. falciparum* development within *An. gambiae* and *An. coluzzii*.

## Background

Despite progress in control efforts over the past decade, malaria remains world’s deadliest infectious disease with over 200 million reported cases each year [1]. According to the latest estimates of the World Health Organization, 241 million cases of malaria and approximately 619,000 deaths were recorded in 2021. African Region shoulder the heaviest malaria burden with 234 million malaria cases and 593 000 of deaths with 78.9% of deaths in children aged under five years [2]. The transmission of these malaria-causing parasites to humans is exclusively caused by *Anopheles* mosquitoes of which five species (*An. gambiae*, *An. funestus*, *An. arabiensis*, *An. moucheti* and *An. nili*) have been identified as the major malaria vectors in Africa [3]. The use of insecticides against vector mosquitoes and antimalarial drugs remains the mainstay of current malaria control. Unfortunately, some studies showed that the effectiveness of these measures is threatened by the rapid expansion of insecticide resistance in vector populations [4-6]. Clearly, additional strategies are needed in order to combat this deadly disease. The possibility to develop new mosquito control strategies that either suppress (reduction or elimination of the wild mosquito population) or replace with mosquitoes that are unable to transmit pathogens is hugely desirable.

The use of microbial symbionts to reduce the competence of vectors involved in pathogen transmission has gained much attention as an emerging alternative approach towards disease control [7, 8]. One strategy termed paratransgenesis exploiting symbiotic bacteria hasbeen proposed as tool in fighting malaria acting mainly by killing *Plasmodium* ookinete/oocysts in the mosquito midgut [9, 10]. Several symbiotic bacterial species isolated from mosquitoes have been explored as potential paratransgenesis candidates for malaria. For example, *Pantoea agglomerans* and *Serratia marcescens AS1*, are able to propagate through mosquito populations and can significantly reduce *P. falciparum* development when engineered to secrete antiplasmodials [11, 12]. However, *P. agglomeran*s has no drive mechanism to spread throughout mosquito populations, while *Serratia* sp. are major opportunistic pathogens in humans, causing many nosocomial infections every year [13] and have been shown to increase pathogens in other mosquito systems [14].

*Asaia* is a promising candidate that colonizes the ovaries, testes, salivary glands, and the midgut of the mosquito, and spreads both horizontally and vertically through mosquito populations [15-18]. This bacterium colonizes a number of arthropods, especially those that feed on the nectar of plants, including *Anopheles stephensi*, *An. gambiae, An. funestus, An. maculipennis, Aedes aegypti*, *Ae. albopictus*, *Culex pipiens*, *Scaphoideus titanus*, and *Sogatella furcifera*, most of which can vector human diseases [15, 18-24]. Importantly, bacteria in the genus *Asaia* apparently cause few human infections. The presence of *Asaia* in various species of *Anopheles* could be beneficial and could be an applied step toward achieving the applying of paratransgenesis against malaria. Therefore, the high *Asaia* prevalence in these *Anopheles* mosquito species offers a compelling opportunity to study how this endosymbiont bacterium interacts with *Plasmodium* in the mosquito host.

In this study, we investigated the effect of *Asaia* on *P.*L*falciparum* infection in a natural *An.*L*gambiae sl* population in two different eco-geographical localities in Cameroon. We screened mosquitoes to detect the presence of *Asaia spp* and *P.*L*falciparum* using a newly developed qPCR assay. We further quantified the density of both microorganisms in *An.*L*gambiae sl* specimens. With this approach, we empirically assessed the *Asaia sp* effect on *P.*L*falciparum* infection in mosquitoes under natural conditions. Our results contribute to the understanding of the tripartite interactions between microbes, the mosquito host and Plasmodium and its potential value to the development of new strategies for malaria control.

## Methods

### Study sites and Mosquito sampling

Mosquitoes which were used in this study, originated from field and laboratory-reared colonies of adult *Anopheles spp*. Laboratory-mosquitoes were used for bacteria isolation and field mosquitoes were used to identify and quantify *Asaia* bacteria. Field mosquito sampling was done in two localities in Cameroon, namely Gounougou (Northern region 9°03′00″N, 13°43′59″E) and Bankeng (Centre region, 4°40’26.4’’N, 12°22’30’’E.). Wild populations of *An. gambiae* females were collected between 06:00 am and 09:00 am in both study sites during the dry season from December 2018 to January 2019, and during the wet season in August 2019. Mosquitoes were collected using Prokopack electrical aspirators (John W. Hook, Gainesville, FL, USA) following verbal consent from the village chief and each household representative. The collected mosquitoes were kept in individual tubes and subsequently transported to the insectary at the Centre for Research in Infectious Diseases (CRID), Yaoundé for further experiments.

### *Asaia* isolation and morphological identification

Alive *Anopheles* laboratory reared mosquitoes (*An. gambiae* s.l) were anesthetized by incubating them for 2 min at -20 °C and their surfaces were sterilized by washing in 70% ethanol and rinsed in distilled water. The isolation of *Asaia* was then proceeded as described previously by Rami *et al*., 2012 [25] with slight modifications. After sterilization, 5 pools of 5 mosquitoes each were suspended and grinded in 1 ml of sterile saline solution (0.9% NaCl). Then, a 0.5 ml aliquot of this suspension was inoculated into a specific enrichment broth media for isolation of the acetic acid bacteria comprising 2.2% D-sorbitol, 0.5% peptone, 0.5% yeast extract, 100 ppm cycloheximide and its pH was adjusted to 3.5 with hydrochloric acid. Next, isolated bacteria were inoculated in 10.0 ml of handmade enrichment culture medium in culture tubes. When microbial growth occurred, the extent of the growth was determined by turbidity monitoring. After that the suspension of the microorganisms was rubbed on a CaCO3 agar plate which contained 2.2% D-glucose, 1.0% ethanol, 1.0% yeast extract, 0.7% CaCO3 and 1.2% agar. *Asaia* colonies were experimentally identified by the morphological properties and formation of carbonate dissolution haloes in agar plates. All the colonies which could clear the CaCO3 were selected and isolated to perform more experiments.

### DNA extraction from bacterial isolates, field mosquitoes and mosquito species identification

Prior to genomic DNA extraction, individual adult female mosquitoes were surface sterilized by washing them in 70% ethanol for 5 minutes and then rinsing twice with sterile distilled water in order to remove superficial bacteria to avoid external contamination. DNA was extracted from whole individual mosquitoes and bacteria isolates using Genejet extraction kit following the manufacturer’s recommendations. DNA extracts were eluted in a final volume of 100 μl and stored at -20°C for further analysis. Molecular species identification was undertaken using diagnostic species-specific PCR assays to distinguish between the morphologically indistinguishable sibling mosquito species of the *An. gambiae s.l.* complex using the Short Interspersed Elements (SINE) [26].

### Real-time PCR for the detection and quantification of *Asaia* in field mosquitoes

*Asaia* detection targeting the 16S *rRNA* gene using primers AsaH1 and AsaR (Table S1) designed in this study from the sequence obtained through 16sRNA Illumina sequencing [27] was undertaken on a subset of samples collected. qPCR reactions were prepared using 5 μl of FastStart SYBR Green Master mix, a final concentration of 1 µM of each primer, 1 μl of PCR grade water and 2 μl template DNA, to a final reaction volume of 10 μl. Prepared reactions were run on a Stratagene MX3005P (Agilent Technologies, Santa Clara, California, USA). System for 15 minutes at 95°C, followed by 40 cycles of 95°C for 15 seconds and 58°C for 30 seconds. Amplification was followed by a dissociation curve (95°C for 10 seconds, 65°C for 60 seconds and 97°C for second) to ensure the correct target sequence was being amplified. Cycle threshold (CT) values were based on duplicate samples. Positive controls were a previously cultured *Asaia* infected sample. A standard curve was generated from 10-fold 9 serial dilutions of the cultured *Asaia* DNA (two technical replications for each dilution) corresponding to a range of 1.44X10^5^ to 1.44X10^12^ genome/μL. The bacterial colony copy numbers was calculated using the following formula[28]: Copy number= (DNA amount (ng) × 6.022 × 1023/length (bp) × 109 × 650). All amplification products were quantified and normalized using the standard range achieved with the *Anopheles* ribosomal gDNA purified PCR product using primers RPS7-F and RPS7-R [29]. In order to compare bacteria densities between individual samples, relative ratio was calculated by dividing the amount of *Asaia* DNA obtained by absolute quantification by the amount of the housekeeping DNA (S7) determined in the same sample.

### Sequencing and phylogenetic analysis of bacterial isolates and PCR positives samples

To confirm the bacterial isolates, amplified bacterial isolates products were sent for Sanger sequencing. In addition, in order to investigate whether the strains of *Asaia* infecting field mosquitoes differ from one season to another in both localities, samples from Bankeng and Gounougou were subjected to PCR as previously described [18] and also send for sequencing. Overall, four colonies of isolates and 57 individuals field samples were analysed using 16S rRNA gene. Among these 57 samples, the distribution included: 19 and 20 samples from Bankeng dry and rainy season respectively; 5 and 13 samples from Gounougou dry and rainy season respectively. The number of samples used was based on the number of infected in the various species and the clarity of the sequences.

PCR products of *Asaia* positive samples were cleaned up using Exonuclease I (Exo I) and Shrimp Alkaline Phosphate (Exo-SAP protocol) according to the protocol of New England Biolabs (NEB, MA, and USA) and sent for sequencing. Maximum Likelihood phylogenetic trees were constructed using the *Asaia*-specific 16S rRNA gene target. The taxonomic relationships of the strains obtained from this study were inferred against Genbank sequence isolates. These sequences were aligned with ClustalW multisequence alignment tool in BioEdit software with previous existing isolates. *Neoasaia chiangmaiensis* (FJ887939.1), *Gluconobacter oxydans* (KU255083.1) and *Acetobacter tropicalis* (JF930138.1) were used as outgroups. The evolutionary history was inferred by the Maximum Likelihood method using MEGA X based on the Jukes-Cantor model. The robustness of the individual branches was estimated by bootstrapping with 1000 replicates.

### Real-time PCR for the detection and quantification of *Plasmodium falciparum* in mosquitoes

The detection of *Plasmodium* infection from each whole individual mosquito was performed using TaqMan assay as previously described [30]. Absolute quantification of all positives specimen was done using the standard curve using 3D7 strain DNA were generated from serial dilution methods and resulting in a quantification range of 7.10^1^ to 7. 10^5^ genomes/ μL. All amplification products were quantified and normalized using the standard range achieved with the S7 gDNA purified PCR product. In order to compare parasite densities between individual samples, the relative ratio was calculated by dividing the amount of *P. falciparum* DNA obtained by absolute quantification by the amount of the housekeeping DNA (S7) determined in the same sample.

Finally, the analysis of the data was performed using Graph-prism software. The prevalence of *Asaia* and *Plasmodium* were compared using Chi-2 test while intensities of infection were compared using the non-parametric Mann-Whitney test.

## Results

### Morphological and molecular characterization of *Asaia* isolates

To generate the positive control for our real-time PCR, *Asaia* bacteria were isolated from five pools of mosquitoes (*An. gambiae* s.l). Four types of colonies were obtained after culturing with colony ranging in colors (Fig. 1A). PCR analysis confirmed the presence of *Asaia* spp. in all the bacterial isolates. Analysis of the sequences obtained from those isolates with nucleotide BLAST revealed that they were related to the 16S rRNA gene of *Asaia*. These sequences were submitted to the GenBank database with the accession numbers ON54736-ON54755. The constructed phylogenetic tree showed that our isolated samples were completely similar and located in the same cluster (Fig. 1B).

**Figure 1:**
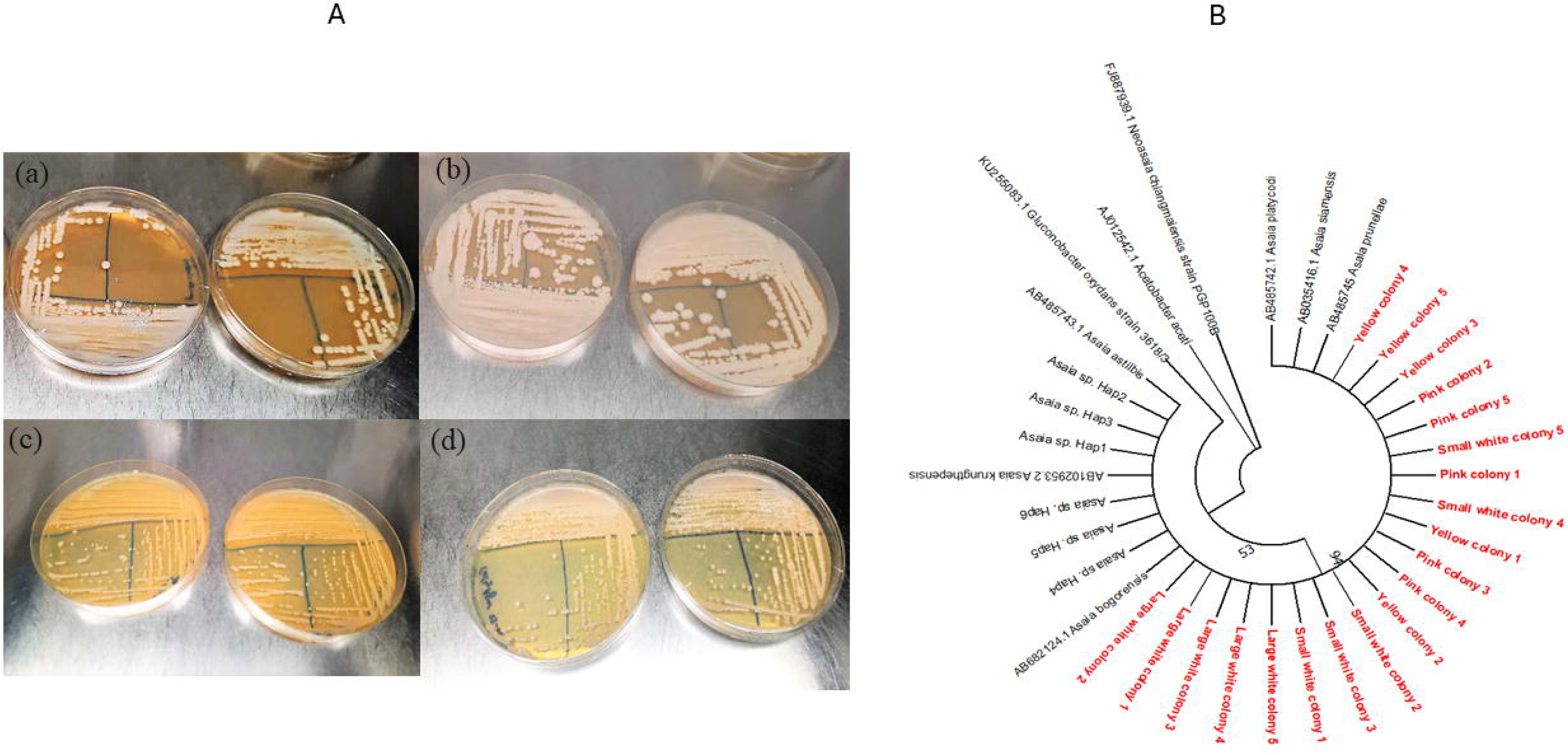
(A) The appearance of *Asaia* colonies on the glucose-yeast extract-CaCO3 medium. Four types of colonies were obtained after culturing with colony ranging in colors from pink to yellowish, white and smooth. *Asaia* species produce white medium non-invasive (a), white large invasive (b), yellow smooth (c) and power pink (d) colonies on glucose-yeast extract-CaCO3 medium which is one of their unique morphological characteristics. (B) Evolutionary analysis of the isolated bacteria. Phylogenetic tree was created with MEGAX software and ClustalW program based on 16S rRNA gene sequences. The Maximum likelihood method was used for evolution relationship analysis and associated taxa were clustered together in the bootstrap test (1000 replicates).

### Mosquito identification and *Plasmodium* spp infection rate

Overall, 1985 mosquitoes were collected including 975 from Bankeng and 1010 from Gounougou. A total of 425 were collected during the dry season and 1560 in rainy season. Molecular identification was conducted on all the 425 mosquitoes collected during the dry season, and 669 randomly selected mosquitoes from the rainy season collection corresponding to 335 and 334 for Bankeng and Gounougou respectively. A predominance of *An. gambiae ss* (n=555, 99%) at Bankeng, during both seasons, with only three *An. coluzzii* individuals. In contrary at Gounougou, the collection was comprised of predominantly *An. coluzzii* (n=503, 93%) and a smaller proportion of *An. arabiensis* 7% (n=36). A subset of 1058 of mosquitoes from Bankeng (n=555) and Gounougou (n=503) were tested for *Plasmodium* infection using a TaqMan assay. The overall prevalence of *P. falciparum* was 16.26 %, with a *P. falciparum* prevalence of 20.1% (112/555) at Bankeng and 11.9% (60/503) at Gounougou. A higher prevalence of *P. falciparum* was observed in *An. gambiae* than in *An. coluzzii*.

### Prevalence, intensity of *Asaia* infection and strain diversity according to the season

A subset of 864 mosquitoes were tested by quantitative PCR for *Asaia,* including 439 and 425 specimens in Bankeng and Gounougou respectively. In Bankeng where only *An. gambiae* ss was identified, the prevalence of *Asaia* was 88.3% while it was 80.9% among *An. coluzzii* mosquitoes in Gounougou (Table 1). A comparison of the medians of infection densities showed that *An. gambiae* mosquitoes (Bankeng) had a significantly higher bacterial load than *An. coluzzii* mosquitoes (Gounougou) (P<0.0001) (Fig. 2A). There was also a significant difference between the infection densities of *Asaia* between seasons in the two localities. Mosquitoes collected in the rainy season (median of 13590) had a higher bacterial density in *Asaia* than those collected in the dry season in Bankeng (median of 64.49) (P<0.0001) (Fig. 2B). In contrast, bacterial densities were significantly higher in the dry season than in the rainy season (P<0.0001) in Gounougou (Fig. 2C). The prevalence of *Asaia* according to the seasons and localities was presented in Table 2.

**Figure 2:**
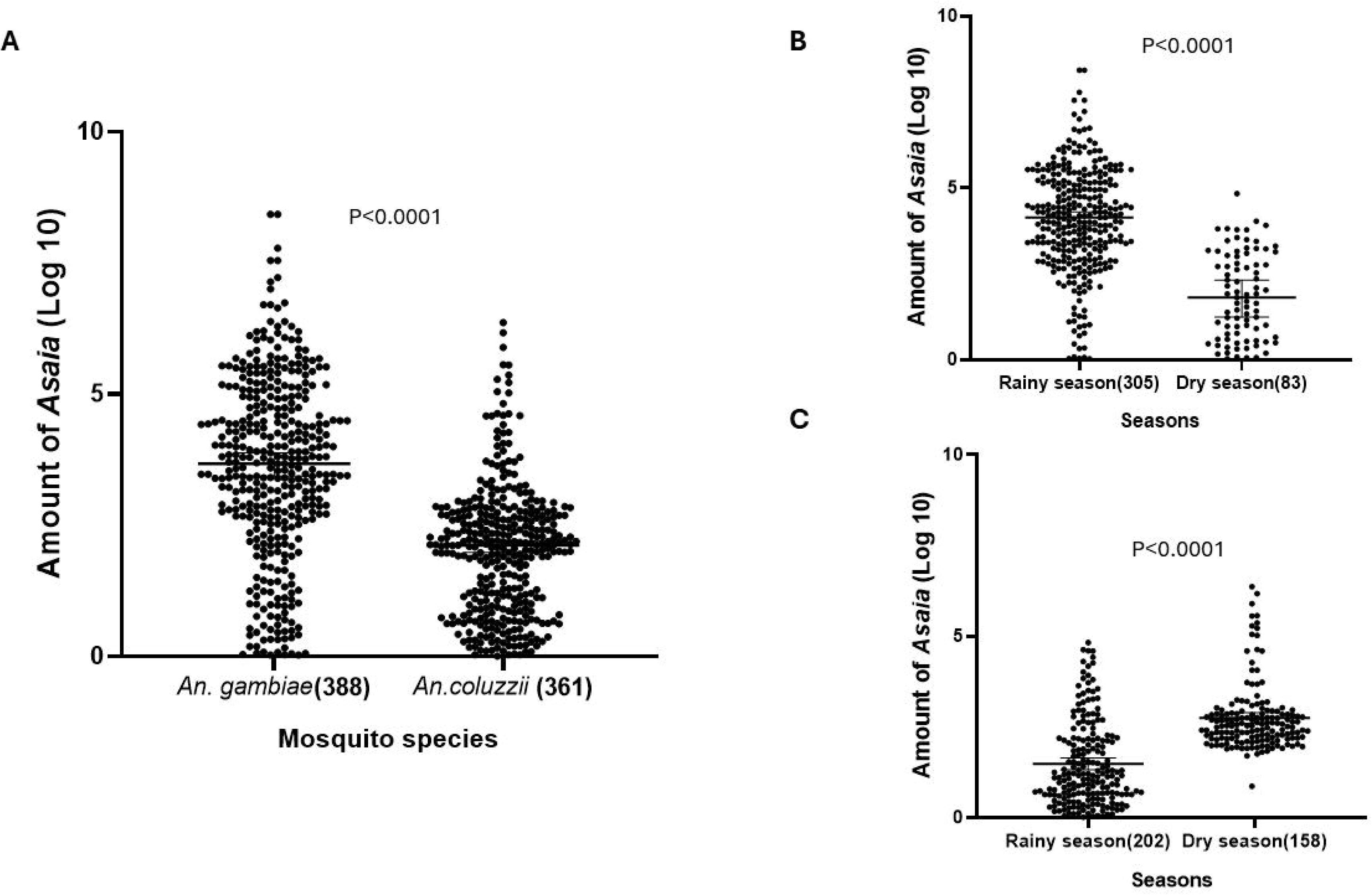
*Asaia* density in Anopheles mosquitoes. Relative amount of *Asaia* in *An. gambiae* at Bankeng and *An. coulzzii* at Gounougou (A). Relative amount of *Asaia* in *An. gambiae* at Bankeng (B) and *An. coluzzii* at Gounougou (C) during Dry and Rainy season. The relative amount of *Asaia* was expressed as a ratio of *Asaia* and mosquito rps7 gene copies in Log10 values; amounts were median with 95% CI. The numbers in the brackets represented the number of samples used for the analysis.

**Table 1:** Prevalence of *Asaia* according to localities or mosquito’s species.

| Localities | *Asaia* positive (%) | *Asaia* negative (%) | Total |
| --- | --- | --- | --- |
| Bankeng (*An. gambiae*) | 388 (88.38%) | 51 (11.62%) | 439 |
| Gounougou (*An. coluzzii*) | 361 (80.94%) | 64 (19.05%) | 425 |
| **Overall** | **749 (86.7%)** | **115 (13.3%)** | **864** |

**Table 2:** Prevalence of *Asaia* according to localities and seasons.

| Localities | Seasons | *Asaia* positive (%) | *Asaia* negative(%) | Total |
| --- | --- | --- | --- | --- |
| Bankeng | Rainy season | 305 (97.7%) | 7 (2.3%) | 312 |
|  | Dry season | 83 (65.3%) | 44 (34.7%) | 127 |
| Gounougou | Rainy season | 202 (76.3%) | 63 (23.7%) | 265 |
|  | Dry season | 158 (99.4%) | 1 (0.6%) | 159 |

Moreover, we have investigated the genetic diversity of *Asaia* sp in *An. gambiae* and *An. coluzzii* according to the seasons. The sequences obtained were subjected to BLAST search in GenBank (NCBI) and all had homology with *Asaia* sp. The relationship between *Asaia* sp. and *Anopheles* mosquitoes revealed clustering of *Asaia* strains in both localities irrespective to the season (Fig. 3A) suggesting that season does not have an impact on the distribution of *Asaia* strains. By analysing the haplotype network, we found eight distinct nucleotide sequence variants (Fig. 3B) where the major one H1 and the others H2, H3, H3, H4 and H5 are present only at Bankeng. In addition, the nucleotide sequence variants H6, H7 and H8 are present only in Gounougou and H8 occurs only in the rainy season. These results can suggest that the variants are localities dependent.

**Figure 3:**
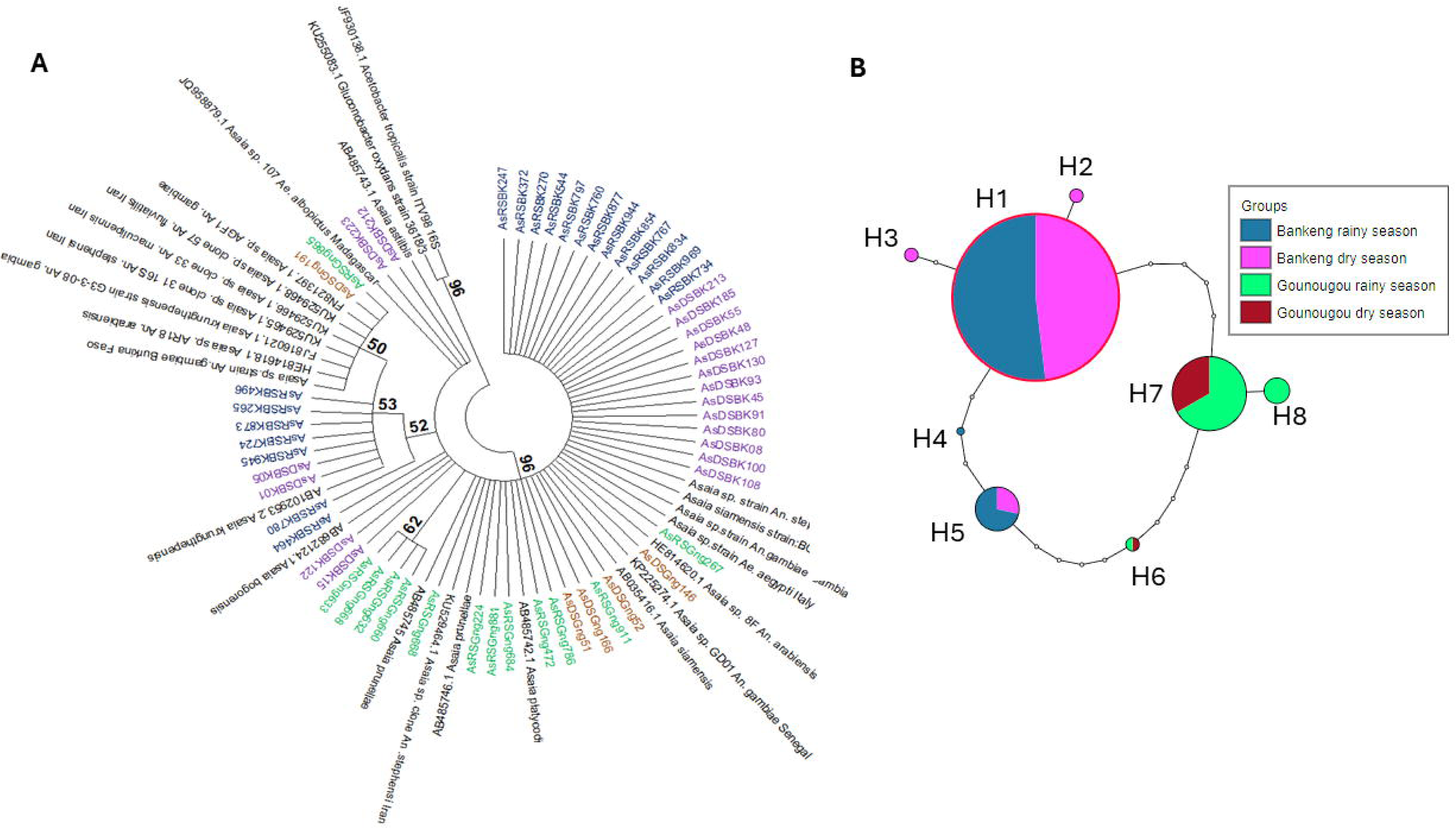
(A) Phylogenetic tree of the strains of *Asaia* in both seasons in both localities. AsRSBK represent samples of rainy season at Bankeng (*An. gambiae*), AsDSBK (purple) samples of dry season at Bankeng; AsDSGng (blue) and AsRSGng (green) represent samples of dry and rainy season respectively at Gounougou (*An. coluzzii*). (B) Nucleotide sequence variant network. Nucleotide sequence variant networks were constructed using *Asaia* sequences in the TCS program. H1–H8 represent the different nucleotide sequence variants, and each colour represents each mosquito species according to locality and season.

### Correlation between *Asaia* sp. and *Plasmodium falciparum* infections

Beyond the spatio-temporal variation of *Asaia* bacteria in *An. coluzzii* and *An. gambiae*, the correlation of *Asaia* infection and *Plasmodium* infection status was also evaluated. The observed proportion of females co-infected with *Asaia* sp. and *Plasmodium* was 13% (58/488) and 14% (59/423) in Bankeng (*An. gambiae*) and Gounougou (*An. coluzzii*) respectively. The prevalence of *Asaia* was 79.45% (58/73) in *Plasmodium*-infected mosquitoes and 90.2% (330/366) in *Plasmodium*-uninfected mosquitoes in Bankeng. In Gounougou, all *Plasmodium*-infected mosquitoes were infected with *Asaia* (100%) while the prevalence of *Asaia* was 82.7% (301/364) in *Plasmodium*-negative mosquitoes (Table 3). The observed proportion of females co-infected with *Asaia* sp. and *Plasmodium* (58/488 = 13%) (Fig. 4A) in Bankeng was similar to expected (14.1%) based on the prevalence of *Plasmodium* (18.8%) and *Asaia* (75.1%) infection (expected prevalence = 0.188× 0.751=0.141) indicating that females infected with one organism are likely to be infected with the other. In Gounougou (*An. coluzzii*), 301/423 (71.1%) females were infected with *Asaia* sp. The observed proportion of females co-infected with *Asaia* sp. and *Plasmodium* (59/423 = 14%) was lower than expected (9.8%) based on the prevalence of *Plasmodium* (13.9%) and *Asaia* (75.1%) infection (expected prevalence = 0.139× 0.711=0.988), indicating that females infected with *Asaia* could facilitate *P. falciparum* infection (Fig. 4B).

**Figure 4:**
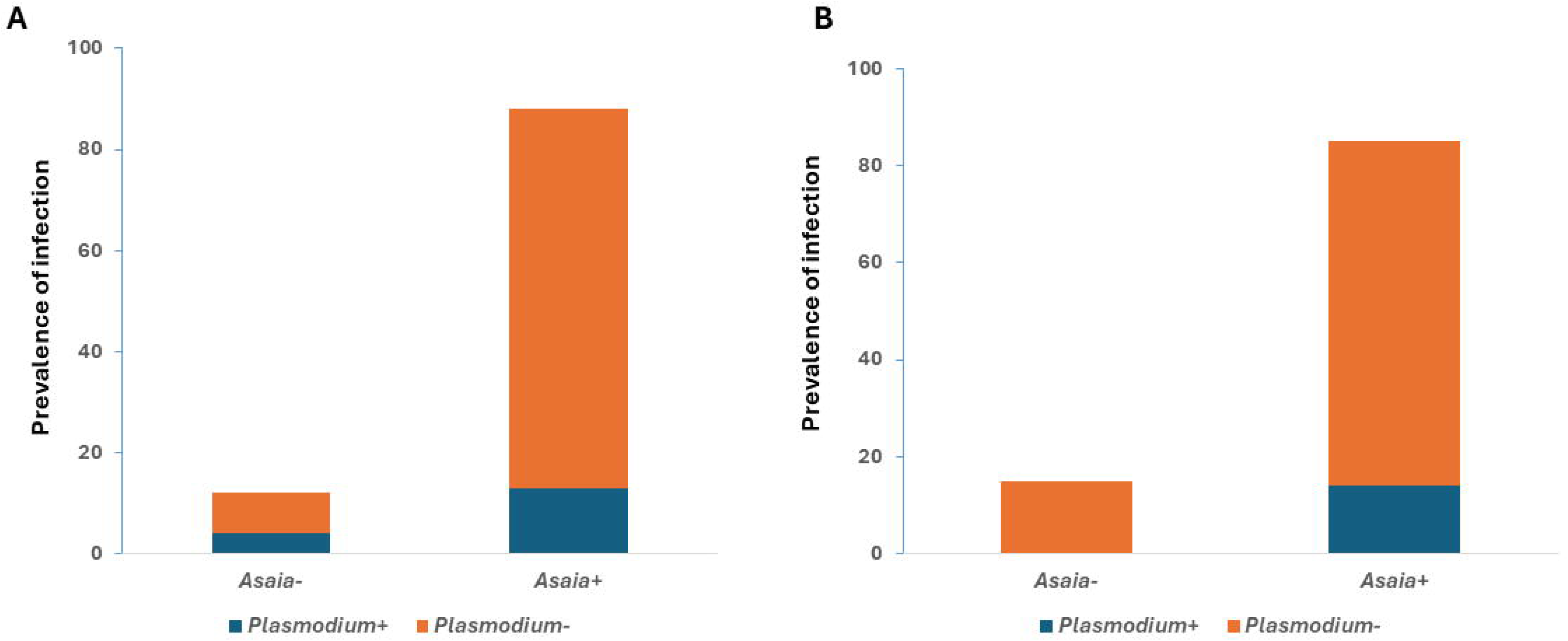
Prevalence of *Asaia* and *Plasmodium* in *An. gambiae* and *An. coluzzii* mosquitoes from Bankeng and Gounougou respectively. A) *Plasmodium falciparum* and *Asaia* infection rates (%) in *An. gambiae* females B) *Plasmodium* and *Asaia* infection rates (%) in *An. coluzzii* females. Add significance sentence.

**Table 3:** Prevalence of *Asaia* according to localities and *Plasmodium* infection status.

| Localities | Infection status | *Asaia* positive (%) | *Asaia* negative (%) | Total |
| --- | --- | --- | --- | --- |
| Bankeng | *Plasmodium*+ | 58 (79.45%) | 15 (20.55%) | 73 |
|  | *Plasmodium*- | 330 (90.2%) | 36 (9.8%) | 366 |
| Gounougou | *Plasmodium*+ | 60 (100%) | 0 | 60 |
|  | *Plasmodium*- | 301 (82,7%) | 63 (17.3%) | 364 |

The median densities of *Asaia* infection were subsequently compared between *Plasmodium*-infected and non-infected mosquitoes. A significant difference between the infection densities of *Asaia* in *Plasmodium* infected and uninfected mosquitoes in the two locations was observed. Specifically, in Bankeng, *Plasmodium*-uninfected *An. gambiae* ss mosquitoes had a higher *Asaia* density compared to *Plasmodium*-infected mosquitoes (P<0.0001) (Fig. 5A). In contrast, *Plasmodium*-infected *An. coluzzii* had higher *Asaia* loads compared to their *Plasmodium*-free counterparts (P<0.0001) (Fig. 5B).

**Figure 5:**
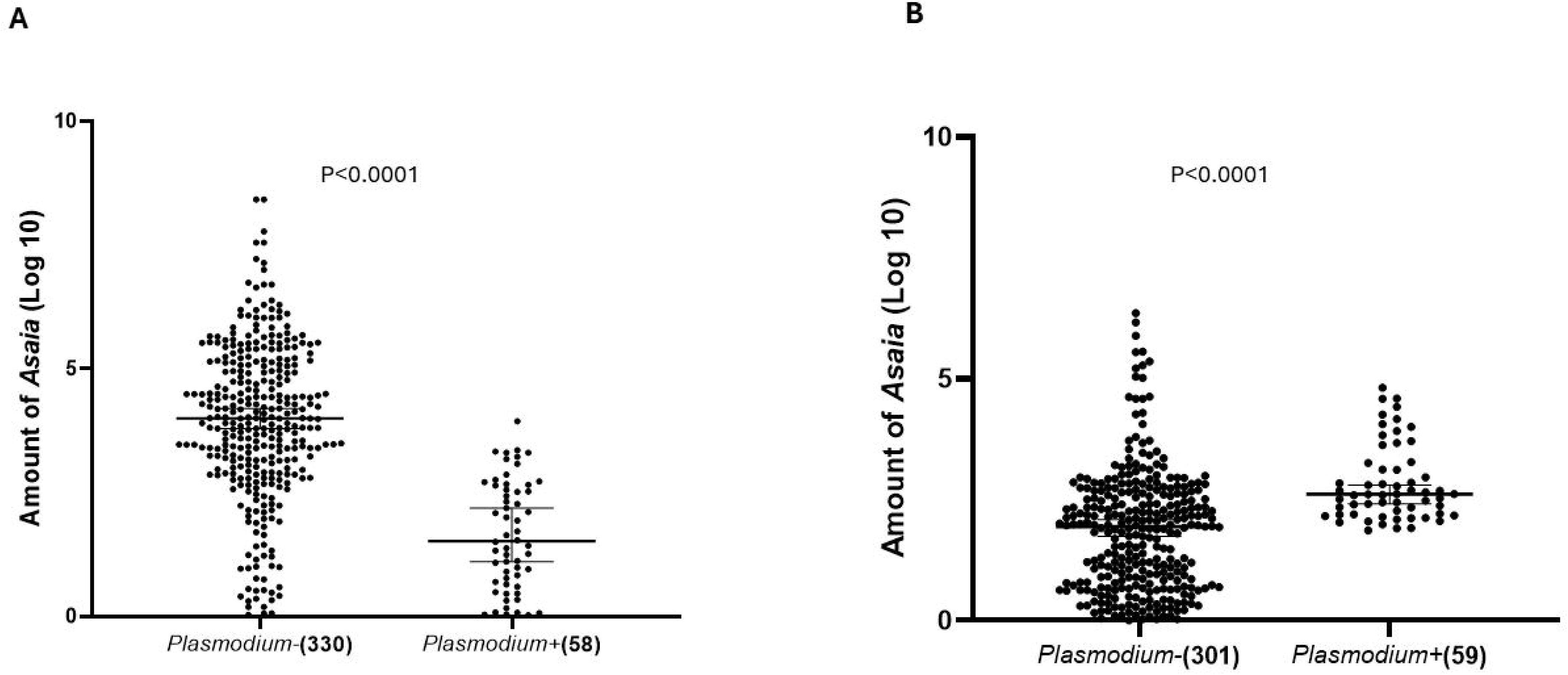
Relative amount of *Asaia* in *Plasmodium* infected and non-infected *An. Gambiae* (A) *and An. coluzzii* (B). The relative amount of *Asaia* was expressed as a ratio of bacterial 16S rRNA and mosquito rps7 gene copies in Log10 values; amounts were median with 95% CI. P. *falciparum*+ represents *Plasmodium falciparum* positive samples and *Plasmodium-*, non-infected *Plasmodium* samples. The numbers in the brackets represented the number of samples used for the analysis.

In order to understand the interaction in more detail, we examined the correlations between *Asaia* and *P. falciparum* densities. Our data showed a very weak positive correlation between *Asaia* and *P. falciparum* (p>0.05) in both localities with similar correlation curves (R^2^=0.02 and R^2^=0.05 for Bankeng and Gounougou respectively (Fig. 6A and Fig. 6B). This result reflects an independent evolution of the two organisms suggesting that *Asaia* and *Plasmodium* do not have an impact on each other in mosquito natural populations studied.

**Figure 6:**
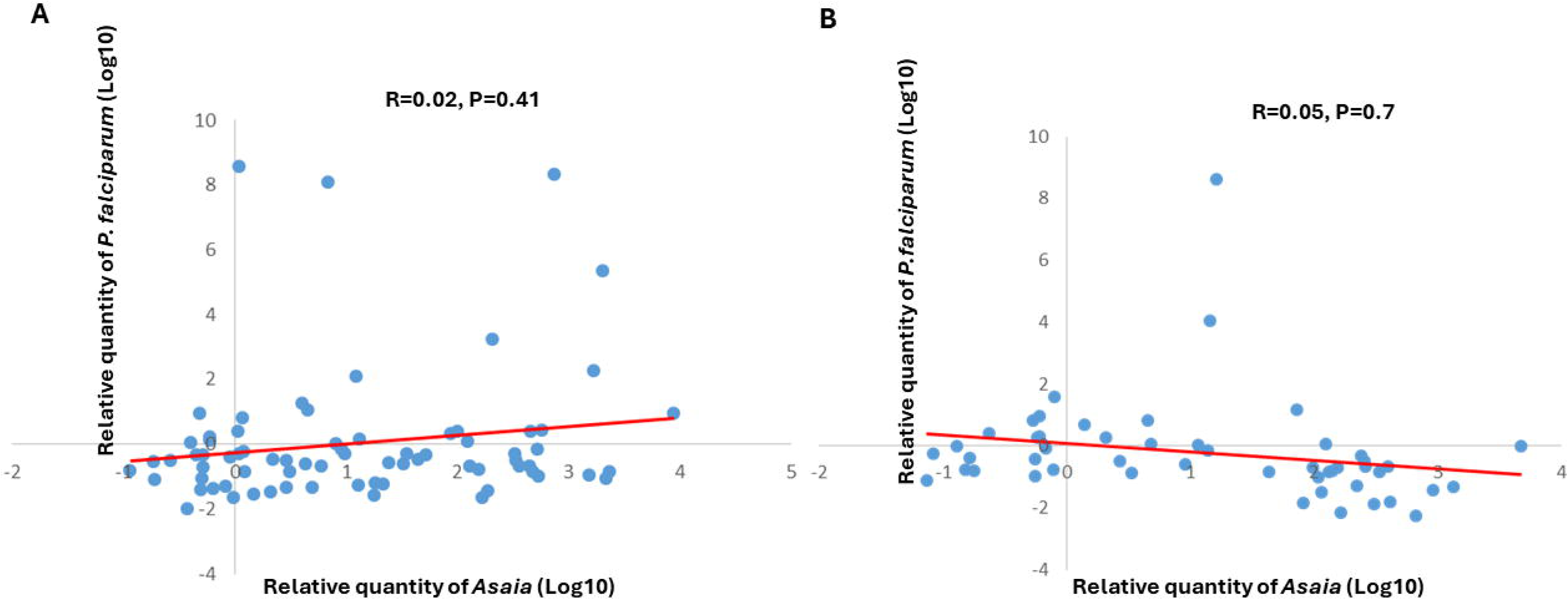
Correlation curves between *Asaia* and *P. falciparum* densities. Relative quantities of *Asaia* obtained from individuals presented and the corresponding *Plasmodium* parasites numbers were reported. (A) Bankeng, (B) Gounougou.

## Discussion

Paratransgenesis is a promising and particularly ingenious strategy currently being developed for controlling vector-transmitted diseases. It utilizes the genetically manipulated native microbiota of the insect vector to inhibit or kill the pathogen within the insect vector. Among the several candidates of paratransgenic control of malaria, *Asaia* is the focus of consideration due to its specific features such as stable association with *Anopheles* mosquitoes, genetic tractability, and horizontal and vertical transmission modes [18]. This study assessed the infection rate and density of *Asaia* sp. and the correlation with *Plasmodium* in *An. gambiae* and *An. coluzzii* in Cameroon.

The *Anopheles* mosquito species studied here have an overall *Asaia* prevalence of 86.7%, with a similar prevalence of *Asaia* sp. between *An. gambiae* and *An. coluzzii* suggesting that both species are susceptible to the infection by *Asaia* sp. This infection rate of *Asaia* is higher than the one obtained in the previous study in *Anopheles* species in Cameroon [18]. This difference in prevalence could be due to seasonal and geographical heterogeneities variation in microbiota composition in *Anopheles* mosquitoes from the various studies sites [31-33]. These contrasting results can also suggest that environmental conditions in which mosquito larvae develop are key determinants within the vectors and this could have important implications on the composition of the microbiota of mosquitoes [33]. Finally, this difference could be also explained by the methods of detection used; in fact, in this latter the detection was done by conventional PCR while in the present study, we used quantitative PCR which is a more sensitive technique [34]. We then quantified *Asaia* density in its mosquito host and investigated variations of these densities according to season and localities. We first observed that *Asaia* densities varied greatly in all our individual samples. The analysis of the densities of infection showed that *An. gambiae* are more highly infected than *An. coluzzii*. In addition, the results showed a higher density infection of *Asaia* in the dry season compared to the rainy season for *An. coluzzii,* meanwhile *An. gambiae* mosquitoes collected in rainy season were more highly infected by *Asaia* than those collected during dry season. Previous study revealed that the larval habitats of *An. gambiae* s.l., may harbor diverse microbial communities and the composition may differ due to microbial interaction [35]. The authors showed that genome comparison of *An. gambiae* and *An. coluzzii* revealed that immune genes are highly diverged [35]. They hypothesized that divergence is driven by an immune response to microbes specific to the different larval aquatic habitats occupied by these species [35]. In addition this difference in densities of infection can be also due to host age and the physiological status can greatly influence its density, as usually observed in other mosquitoes species [36, 37]. Alternatively, microbe-microbe interactions can influence prevalence and density of members of the microbiome, and effect the host [38, 39] and *Asaia* is known to antagonize and interact with other microbes [24, 40].Our *Anopheles* mosquitoes were from a natural population, therefore, age or physiological cycle were unknown, which may explain the variation of *Asaia* densities we observed, which could affect *Plasmodium* infection.

*Asaia* has been shown to be a promising candidate for paratransgenic control of malaria. This is as a result of its negative effect on the development of *Plasmodium* at different stages [41] and its ability to transmit and persist in field *Anopheles* species [18]. In this study, we focused on the interaction *Asaia*-*Plasmodium* in natural *Anopheles gambiae* s.l populations by quantifying the amount of *Asaia* in *Plasmodium* infected and non-infected mosquitoes. In *An. gambiae*, *P. falciparum* infected mosquitoes had higher densities of *Asaia* compared to *Plasmodium* non-infected suggesting that the presence of *Asaia* can impede the infection by *Plasmodium.* The inhibitory effects of this symbiotic bacteria on *Plasmodium* parasites have been already shown [16, 42]. In fact, *Asaia* has shown to act as an immunomodulator within the mosquito by inducing the production of antimicrobial peptides that may interfere with the course of infection, particularly during the invasion of epithelial tissues and translocation to the salivary glands. *Asaia* may also act as up-regulators of immunity genes [16]. In contrary, in *An. coluzzii* all *Plasmodium*-infected mosquitoes were infected with *Asaia* while the prevalence of *Asaia* was 82.7% in *Plasmodium*-negative mosquitoes, indicating that *Asaia* could facilitate *P. falciparum* infection. Unfortunately, there are a lack of studies on *Asaia* showing the same pattern, making impossible the comparison between other studies. However, another study showed that different strains of the bacteria *Serratia* can induce different outcomes on *Plasmodium* infections [43]. Bai et al identified two mosquito symbiotic *Serratia* strains *Serratia* Y1 and *Serratia* J1, from field caught *An. sinensis* mosquitoes, with different contrasting effect on *Plasmodium berghei* development where *Serratia* Y1 shows anti-*Plasmodium* activity, but *Serratia* J1 does not influence parasite development in the midgut of mosquitoes [44]. These indicate that the effect of mosquito gut bacteria on parasite infection is complex and may depend on species-specific or strain-specific interactions [43]. It was also suggested that despite the fact that there is overlay between the mosquito antibacterial and anti-malarial immune responses, some immune genes induced by bacteria have minimal impact on *Plasmodium* development [45-47]. In *Anopheles* mosquitoes, ingestion of a blood meal containing high loads of *Plasmodium* triggers a strong immune response, acting both in the lumen and in the gut wall [48]. Capone et al [16] also explained that *Plasmodium*-infected blood meals induce an immune response in mosquitoes that might limit not only the malaria parasite progression in the insect but also the symbiont population. The midgut microbiota of the *Anopheles* can be negatively affected by this response [16]. Moreover, the lower level of *Plasmodium* infection observed previously in *An. coluzzii* could be explained by this higher intensity suggesting that *Asaia* influence *Plasmodium* transmission. Indeed, it has been stated that the composition of the gut microbiota may influence parasite transmission and can explain the difference in infection levels between mosquito populations from diverse environments [49, 50]. If the presence of *Plasmodium* reduces the load of symbionts in the midgut of a given mosquito, this would limit the usefulness of the symbiont itself for paratransgenic applications. Further investigations are then needed to better clarify the *An. coluzzii*-*Asaia*-*Plasmodium* system in the localities studied.

## Conclusion

Here we isolated and quantified *Asaia* in *An. gambiae* and *An. coluzzii*, and characterized the *Asaia*-*Plasmodium* interactions in these Anopheline species. We have shown that, seasons, species of mosquito and/or localities impact the amount of *Asaia* and moreover the complexity of the *Asaia*-*Plasmodium* relationship. However, no correlation was observed between the amount of *Asaia* and *P. falciparum* infection, suggesting *Asaia* could be exploited for paratransgenic control. Further studies such as experimental infection following by the immunological experiments are needed to better understand the mechanisms involved in order to clearly characterize *Anopheles*-*Asaia*-*Plasmodium* interactions.

## Availability of data and materials

The dataset supporting the conclusions of this article is available in GenBank database database under the accession numbers ON54736-ON54755.

## Authors Contributions

MMS and CSW conceived and designed the study. MMS and CGM, collected the samples on the field. CGM, MMS and FMD performed the culture and isolation of *Asaia*. CGM, MMS and MT designed the experiments and performed the validation of the qPCR assay. CGM performed the data analysis with the help of MMS. CGM and MMS wrote the manuscript with contribution from, FN, GLH and CSW.

